# Colonization with heterologous bacteria reprograms a *Caenorhabditis elegans* nutritional phenotype

**DOI:** 10.1101/2020.03.01.972349

**Authors:** Qing Sun, Nicole M. Vega, Bernardo Cervantes, Christopher P. Mancuso, Ning Mao, Megan Taylor, James J. Collins, Ahmad S. Khalil, Jeff Gore, Timothy K. Lu

## Abstract

Animals rely on the gut microbiome to process complex food compounds that the host cannot digest and to synthesize nutrients that the host cannot produce. New systems are needed to study how the expanded metabolic capacity provided by the gut microbiome impacts the nutritional status and health of the host. Here we colonized the nematode *Caenorhabditis elegans* gut with cellulolytic bacteria that enabled *C. elegans* to utilize cellulose, an otherwise indigestible substrate, as a carbon source. The nutritional benefits of colonization with cellulolytic bacteria were assayed directly, by incorporation of isotopic biomass, and indirectly, as host larval yield resulting from glucose release in the gut. As a community component in the worm gut, cellulolytic bacteria can also support additional bacterial species with specialized roles, which we demonstrate by using *Lactobacillus* to protect against *Salmonella* infection. As a model system, *C. elegans* colonized with cellulolytic bacteria can be used to study microbiome-host interactions. Engineered microbiome communities may provide host organisms with novel functions, such as the ability to use more complex nutrient sources and to fight against pathogen infections.

**One Sentence Summary:** Heterologous bacteria colonizing an animal gut help digest complex sugars to provide nutrition for the host in a model system.

---

The gut microbiome is a complex community of microbes, the ecology of which is intimately tied to the physiology of its host organism (*1, 2*). These symbiotic microbes have been shown to contribute to the nutritional status and metabolism of the host, train the immune system, and control brain development and behaviors (*3*). Carbohydrates are important sources of energy for both microbes and animals (*4*). Simple sugars like glucose and lactose can be easily absorbed by hosts, but more complex carbohydrates and plant polysaccharides, including cellulose, xylans, resistant starch, and inulin, are not as easily digested. The gut microbiome can confer upon its host, such as termites (*5*) and ruminants (*6*), caloric benefits by breaking down these ingested plant carbohydrates, which the host enzymes cannot digest (*7*). The gut microbiome can also evolve to adapt to new carbohydrate sources when hosts change their eating habits. For example, the Japanese, as a result of their diet, have intestinal microbiomes that have gained algal carbohydrate processing enzymes for seaweed processing through horizontal gene transfer from marine bacteria (*8*). However, research into the use of non-native bacteria to perform nutritional functions for other animal hosts has been limited. An understanding of how the various species of microorganism constituting the microbiome affect their host is fundamental to deriving the potential benefits of manipulating the microbiome.

In this study, we assembled a functional microbiome in a simple animal gut to allow host utilization of a complex carbohydrate source. We used the nematode *Caenorhabditis elegans* as the animal host because it has been extensively used as a model system to elucidate mechanisms of interaction between prokaryotes and their hosts (*9, 10*). Specifically, we colonized *C. elegans* with cellulolytic bacteria that break down cellulose in the gut, such that the released glucose is available as a nutrient for both *C. elegans* itself and the colonizing bacteria. To move this system toward higher community complexity as occurs in a natural gut microbiome, we added an additional bacterial species with the specialized function of preventing infection by pathogens. Overall, our results indicate that by assembling a functional community in the *C. elegans* gut, we can extend the nutrition processing and pathogen inhibition capacities of the animal host.

## Results

To introduce a novel carbon processing capability to *C. elegans* (Fig. 1), we started by selecting cellulose-degrading microbes as potential intestinal colonists. To identify these functional bacteria members, we assembled a collection of *C. elegans* native gut bacteria (*11*) (see Table S1 for a full list) and tested these species for their ability to utilize carboxymethyl cellulose (CMC) and other carbon sources by measuring bacterial optical density following *in vitro* incubation (Fig. S1). Additionally, we checked the cellulose degradation capacity for each strain using a Congo Red assay (*12, 13*). Briefly, after overnight culture on CMC plates and visual confirmation of bacterial growth, we stained the plates with Congo Red. As Congo Red stains glucose polymers, a halo surrounding the colony indicates cellulase activity. None of the native *C. elegans* isolates (Fig. 2A) exhibited detectable cellulase activity, which reflects their lack of cellulose processing capability. To further identify cellulolytic bacteria, we screened a set of soil bacteria (Table S1) and identified *Pseudomonas cellulosa* and *Bacillus subtilis* as cellulase active bacteria (Fig. 2A). Of these strains, *P. cellulosa* produced the highest levels of reducing sugars, including glucose (Fig. 2B), suggesting that this strain could be a potentially useful nutritional endosymbiont for the *C. elegans* host and supply it with nutrients that would otherwise be unavailable.

**Figure 1.**
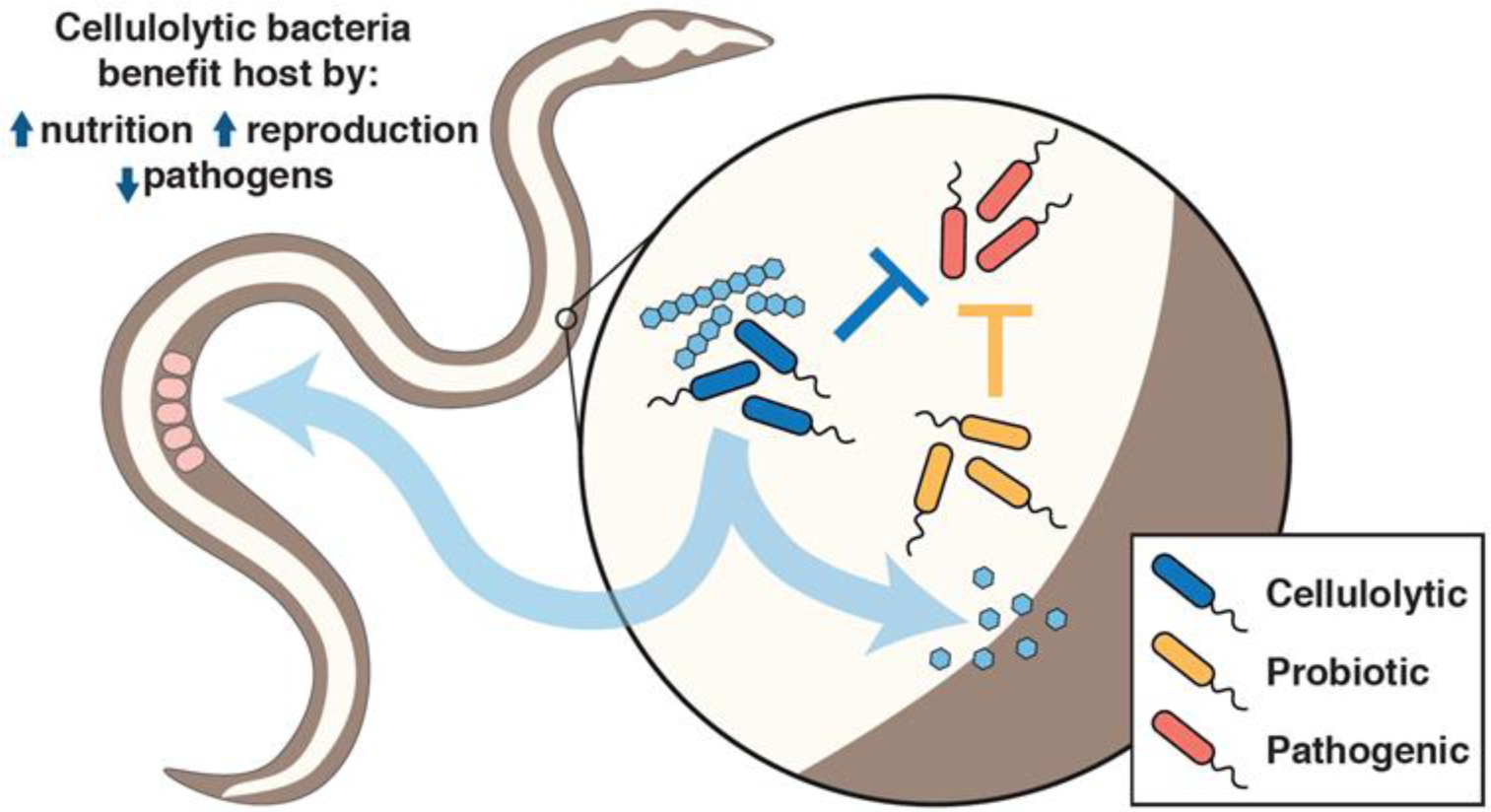
Benefits provided by colonization of the *C. elegans* gut by a heterologous microbial community. Cellulolytic bacteria are able to break down cellulose, such that the released glucose can serve as nutrition for both *C. elegans* and colonized bacteria. Additional bacterial species can improve resistance against pathogenic bacteria.

**Figure 2.**
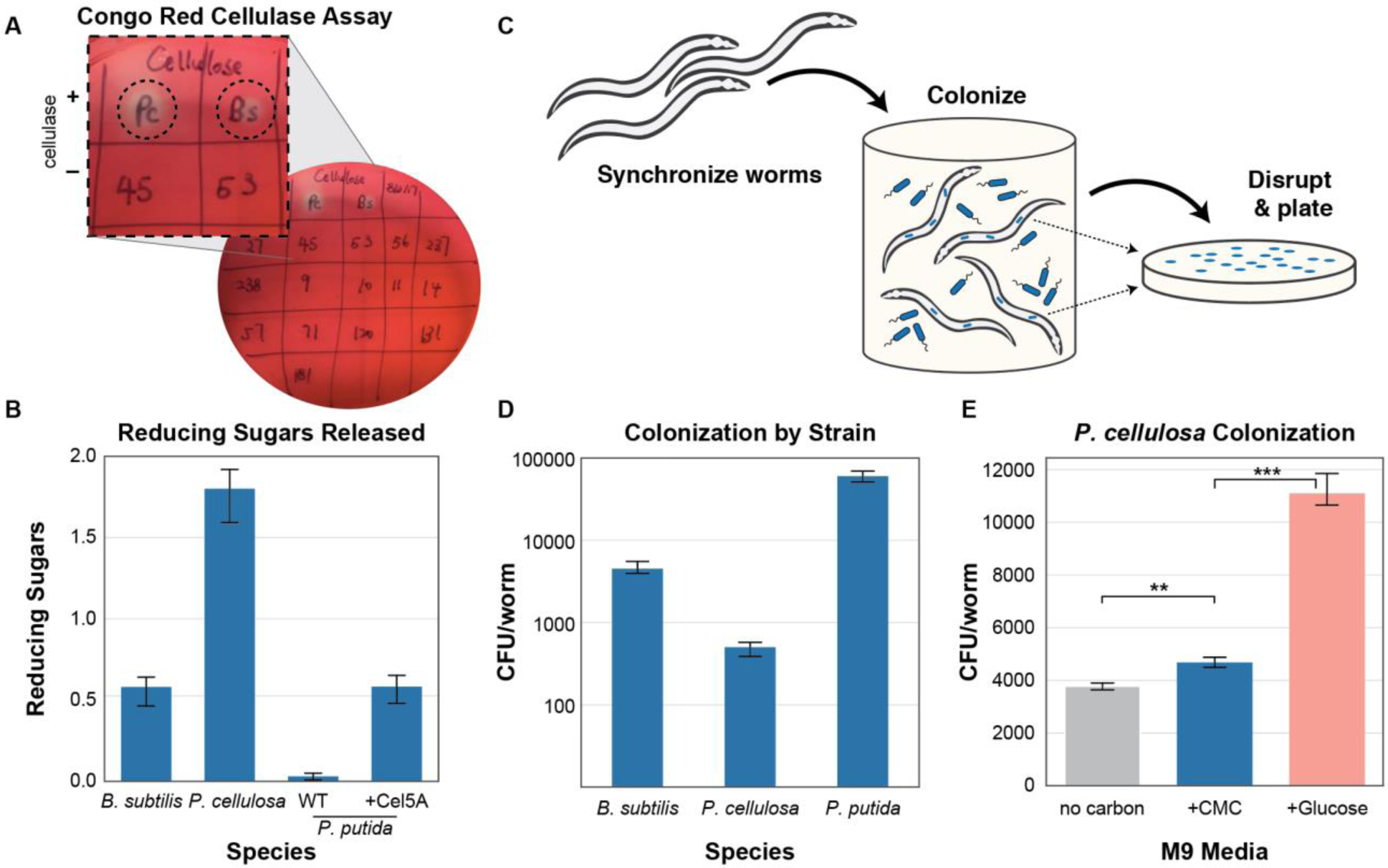
Bacterial colonization and function in worm gut. A) Cellulase functional screening by Congo Red assay. Halos on the plate indicate cellulase activity. Pc: *Pseudomonas cellulosa*, Bs: *Bacillus subtilis.* Numbered strains are from the *C. elegans* native gut microbiome as listed in Table S1. B) Production of reducing sugars by cellulose-degrading bacterial strains. C) Schematic of bacterial colonization and analysis procedure. Synchronized L1 wild-type worms were grown on selected strains of live bacteria and then disrupted for colonization analysis by plating and colony counting. D) Colony-forming units (CFU) of various bacterial species in the worm gut. E) *C. elegans* colonized with cellulolytic *P. cellulosa* and incubated with different carbon sources. **P < 0.01, ***P < 0.001, Student’s t test. Error bars represent 95% confidence intervals of the mean.

We next sought to determine whether a cellulolytic organism could successfully colonize the worm intestine. *C. elegans* feed on bacteria, a certain percentage of which escape digestion and colonize the gut (*14, 15*). For colonization, we allowed synchronized adult N2 worms to feed in liquid bacteria culture to ensure that all the worms experienced a uniform environment (*15*). Worms were then collected, washed, and mechanically disrupted to release intestinal bacteria. These suspensions were then plated on agar to estimate bacterial colonization (Fig. 2C). Naturally cellulolytic bacteria *P. cellulosa* and *B. subtilis*, as well as a natively non-cellulolytic bacterium, *Pseudomonas putida*, which had been engineered to produce endoglucanase (see Methods), achieved a colonization density of ∼10^2^-10^4^ colony forming units (CFU) per worm (Fig. 2D). Colonization was stable for at least two days in the *C. elegans* gut. All of these bacterial species were therefore able to colonize the host intestine, albeit at different population sizes, but at colonization levels comparable to those of native *C. elegans* gut strains (Fig. S2).

To determine whether bacteria identified as being cellulolytic *in vitro* could hydrolyze complex sugars in the gut environment, we pre-colonized *C. elegans* with the cellulolytic strain *P. cellulosa. P. cellulosa* was chosen for its strong ability to degrade CMC and release reducing sugars, including glucose (Fig. 2B). Since *P. cellulosa* grows to low densities in culture, we used *E. coli* OP50 as a supplemental food source to ensure that worms had adequate nutrition during growth. Synchronized N2 worms raised on mixed lawns of *P. cellulosa* and *E. coli* OP50 on solid NGM medium showed intestinal colonization by *P. cellulosa* as young adults 46 hours post-L1 (Fig. 2D), indicating that this organism is able to colonize the host. Pre-colonized worms were then incubated for 24 hours in liquid media (M9 worm buffer) with: 1) no carbon source, 2) cellulose (in the form of CMC), or 3) glucose. After incubation, worms were washed and digested to check gut bacterial density. We found that M9 buffer with CMC as a carbon source enhanced the colonization density of *P. cellulosa* compared with no carbon source (Fig. 2E). This experiment demonstrated that a cellulolytic bacterial strain, *P. cellulose*, in the *C. elegans* gut hydrolyzed CMC and benefited from the released sugar. Glucose as a carbon source further enhanced colonization density because free sugar monomers are an easier-to-access carbon source than CMC.

Next, we sought to determine whether colonization by cellulolytic bacteria would allow the host to use the otherwise indigestible cellulose carbon source. *C. elegans* is able to use simple carbohydrates, including glucose (*16*), but does not produce enzymes that degrade cellulose. We hypothesized that if cellulolytic organisms in the gut could liberate carbon in a form that is available and metabolizable by the host, colonization of the gut with a cellulolytic microbiome would provide this host with the ability to use cellulose as a growth substrate. Again, we chose *P. cellulosa* as the colonizing bacteria. Three approaches were used to investigate whether *C. elegans* utilized glucose released by gut bacteria.

First, we used radionucleotide (^14^C) incorporation as a direct measurement of cellulose-derived carbon incorporation into worm biomass. Briefly, adult wild-type worms were colonized with *E. coli* OP50 (non-cellulose degrader) or *P. cellulosa*, then incubated in S medium with heat-killed *E. coli* OP50 and trace ^14^C labeled CMC (radiolabeled cellulose purified from *Arabidopsis thaliana*) (*17*) for 24 hours to allow bacterial degradation of the substrate to proceed. After incubation on the radionucleotide substrate, worms were washed and treated with an antibiotic cocktail for 24 hours to remove bacteria, in order to eliminate the effects of bacterial biomass on scintillation counts and show the incorporation of cellulose-derived carbon into the germ-free worm biomass. A high-throughput antibiotic susceptibility screen of the bacterial strains (including *C. elegans* native gut bacteria and the soil bacteria in Table S1) was conducted to determine an appropriate antibiotic cocktail to eliminate these species from the *C. elegans* gut (Fig. S3). Clearance by the antibiotic cocktail was tested by imaging worms fed with fluorescently labelled *Pseudomonas putida* (Fig. S4). After clearing the bacterial biomass, worms were mechanically digested in batches (200 worms per sample), and ^14^C incorporation was measured using scintillation counts. ^14^C incorporation was significantly higher for *P. cellulosa* colonized worms than for *E. coli* colonized worms, before and after clearance with antibiotics (Fig. 3A), demonstrating that cellulolytic bacteria in the *C. elegans* gut had hydrolyzed ^14^C labeled CMC and that the hydrolyzed glucose had been taken up by the host, as well as by the microbes.

**Figure 3.**
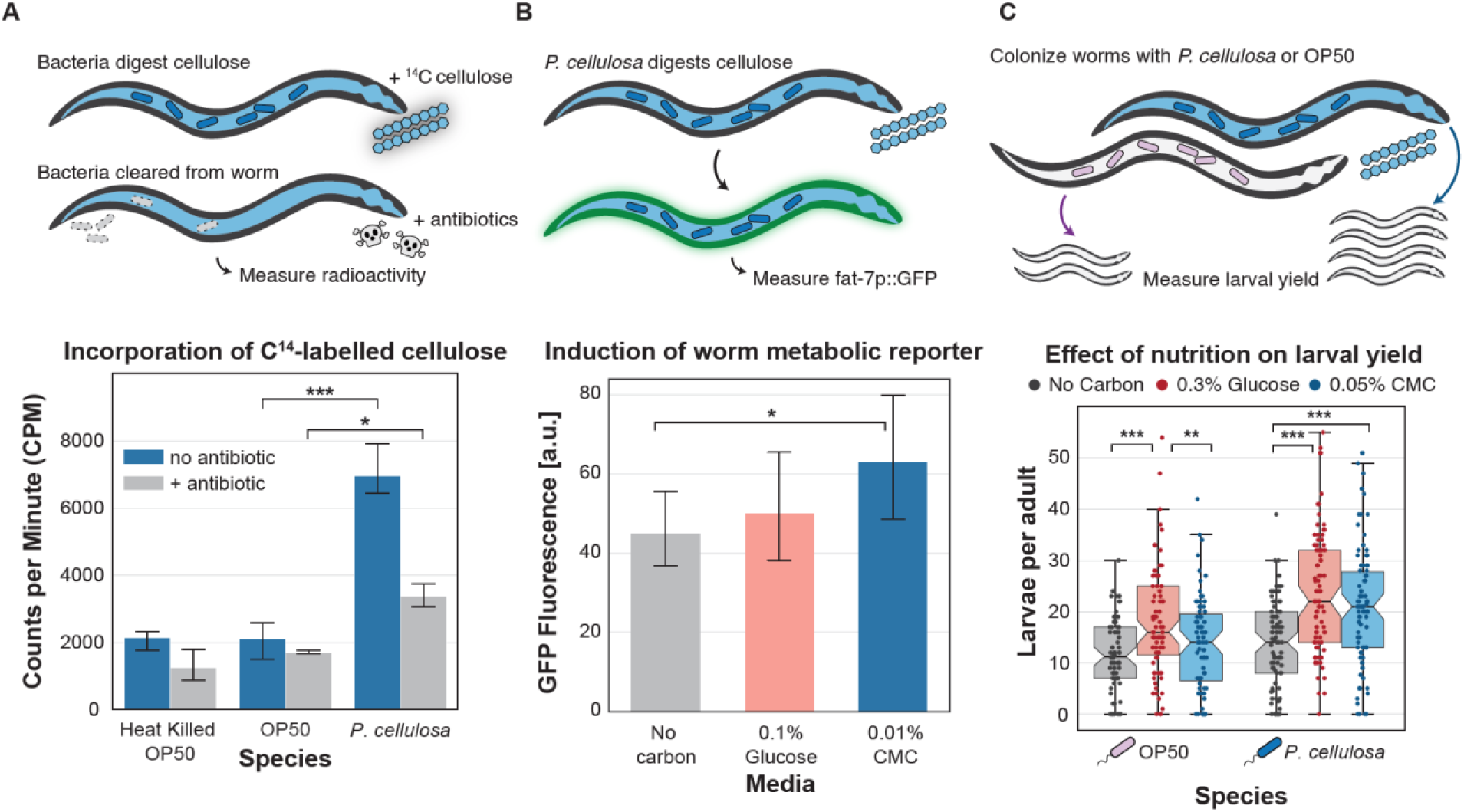
Intestinal cellulolytic bacteria allow host utilization of cellulose. A) Incorporation of ^14^C-labelled cellulose. Worms were first colonized with cellulolytic bacteria and then incubated with ^14^C-cellulose to allow carbon utilization. Antibiotic treatment was applied before ^14^C measurement of worms to eliminate bacterial interference in isotopic reading. ^14^C-cellulose incorporation measurement reflected the isotopic carbon incorporation in *C. elegans*. B) Induction of the worm *fat-7p::GFP* metabolic reporter. Adult worms pre-colonized with *P. cellulosa* were incubated in liquid media for 24 hours with the indicated carbon sources, and GFP fluorescence was read in individual worms on a BioSorter large object sorter. C) Effect of nutrition on larval yield. Pre-colonized worms were separated into the individual wells of a 384-well plate, and larvae counts in each well were read after 48 hours. Larval yield comparison indicates the nutrient benefit of cellulolytic activity. *P < 0.05, **P < 0.01, ***P < 0.001 Student’s t test. In A and B, error bars represent 95% confidence intervals for the mean. In C, notches represent 95% confidence intervals for the median.

Second, we confirmed that incorporation of carbon from microbiome-digested CMC affected host metabolism by using a fluorescent reporter of host nutritional status (*18*). We first confirmed that *fat-7p*::GFP, which is expressed in the intestinal cells of *C. elegans*, was a reliable reporter for carbon uptake by the host, and determined an optimal glucose concentration of 0.1%-0.2% (w/v) for these assays (Fig S5). Young adult wild-type worms (N2 strain, 46 hours after L1 instar) were transferred from plates of live *E. coli* OP50 + *P. cellulosa*, where they had been colonized during growth by live bacteria, to S medium with heat-killed *E. coli* (to provide nitrogen and other nutrients) ± glucose or CMC. Gentamicin (10 µg/mL) was used to prevent the growth of bacteria outside the worms, ensuring that any nutritional benefit would be conferred by resident bacteria in the gut. Worms pre-colonized with *P. cellulosa* showed an increase in fluorescence from this reporter when provided with CMC as a carbon source (Fig 3B), indicating an increase in available nutrients to the host directly due to the microbial liberation of glucose.

Third, we measured the benefits of carbon incorporation into the new host biomass via larval output, to determine whether colonization by cellulose-degrading bacteria could improve the fitness of the host. Larval output in the worm is a function of nutritional status and can be initiated by adding nutrients after starvation at a specific point in the young adult-mature adult transition (*19*). We therefore transferred worms from solid media colonization plates to liquid culture supplemented with glucose or CMC 46 hours after plating L1 larvae, to capture worms at the vulnerable point in this transition. Here, worms were placed into individual wells of a 384-well plate, to allow enumeration of larval yield for single adults. After 48 hours in liquid media with a carbon source, the number of larvae produced by individual worms was counted. Worms colonized with *P. cellulosa*, but not with *E. coli* OP50 alone, produced more larvae per adult worm when CMC was provided in the medium (Fig. 3C, S6), again indicating that colonization by the cellulolytic bacterial species allows the worms to benefit from CMC as a carbon source made available by these resident bacteria.

Alongside these efforts, we sought to determine whether adding a co-colonizing species would produce a community with improved function for the host. As an alternative to adding an additional cellulose-degrading bacterium, we sought to take advantage of the probiotic properties of *Lactobacillus plantarum*, as lactic acid bacteria have previously been shown to have probiotic effects in hosts, including protection against pathogens (*20, 21*). We hypothesized that addition of this probiotic to the gut community could prevent disruption of the community by an intestinal pathogen (here, *Salmonella enterica* LT2). To determine the effect of *L. plantarum* on pathogen colonization, worms were pre-colonized with *P. cellulosa* on solid media as previously described, with and without the addition of *L. plantarum* (10^9^ CFU/plate). After colonization, worms were exposed to *S. enterica* (10^8^ CFU/mL) infection for one hour in liquid culture. Following this initial exposure, worms were washed to remove external pathogens and incubated for 36 hours to allow infection to progress (Fig. 4B). We found that both *P. cellulosa* and *L. plantarum*, alone or in combination, modestly suppressed pathogen proliferation relative to the control condition (colonization by *E. coli* OP50), but there was no significant difference between the *P. cellulosa* only and the *P. cellulosa* and *L. plantarum* combination conditions without a carbon source.

**Figure 4.**
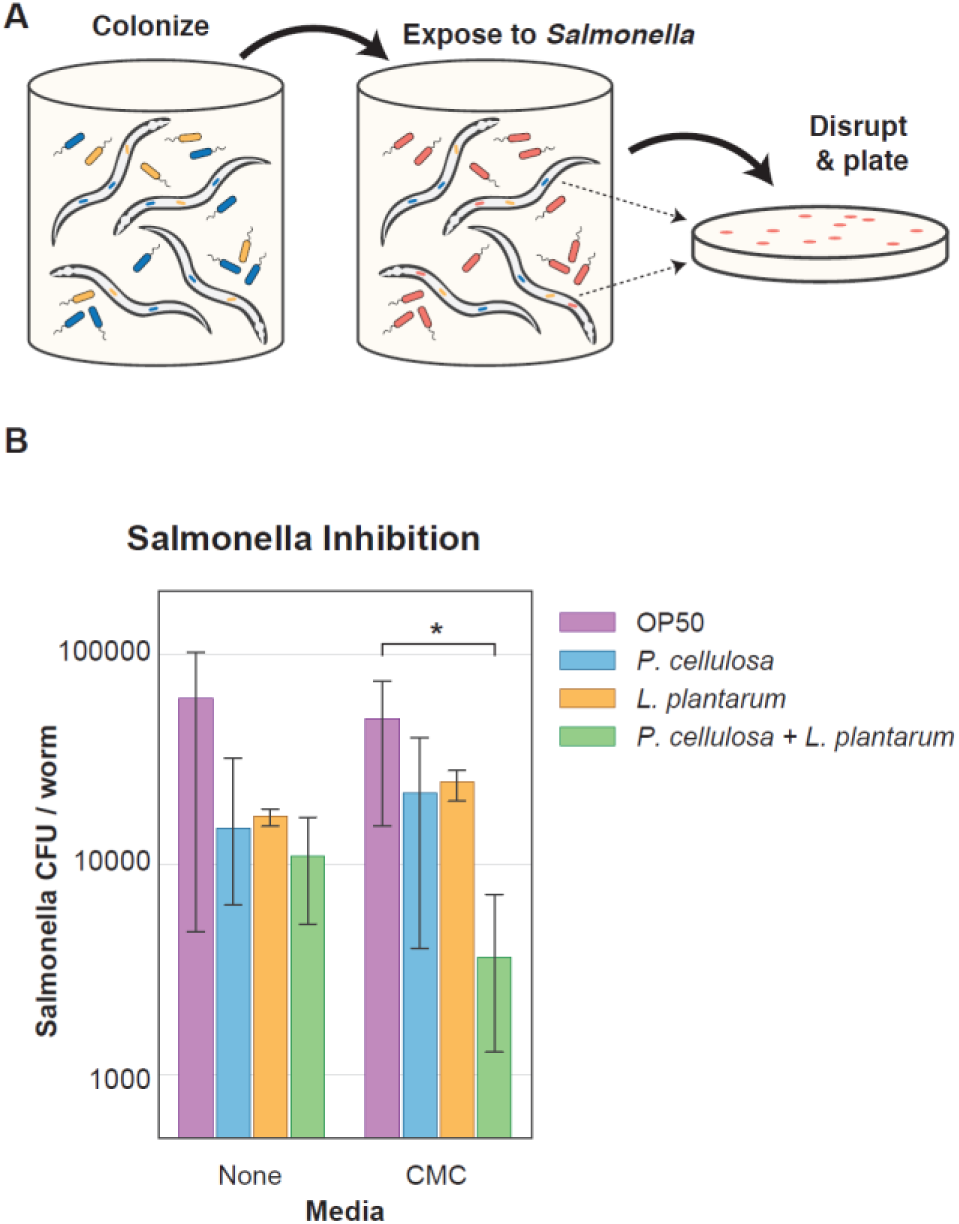
Addition of probiotic *L. plantarum* (Lp) protects *C. elegans* against *Salmonella enterica* (pathogen) invasion of the gut. Synchronized adult worms were colonized for two days on plates containing *E. coli* OP50 + *P. cellulosa* (Pc), *E. coli* OP50 + Lp, or *E. coli* OP50 + Pc + Lp. A) A one-hour exposure to *S. enterica* in liquid culture was used to colonize worms with this pathogen. B) After 36 hours outgrowth of the pathogen, worms pre-colonized with *P. cellulosa* in the presence of *L. plantarum* showed lower pathogen burden than *P. cellulosa* monocolonized worms when CMC was used as the carbon source. Data shown are fold change *Salmonella* infection (CFU/worm) relative to the *P. cellulosa*-only condition. Error bars represent 95% confidence interval of the mean. CMC = 0.01% carboxymethylcellulose (w/v). *P < 0.05 Student’s t test.

In the presence of CMC as carbon source, *P. cellulosa* and *L. plantarum* in combination suppressed the proliferation of *Salmonella* by 10-fold compared with what was seen in worms colonized by a single strain (Fig. 4B). These data suggest that the progression of infection in *C. elegans* could be affected by the resident microbiome. What is more, the presence of CMC as a carbon source enhanced the community’s capability to suppress pathogens, demonstrating that provision of a specific substrate for the cellulose degrader improved overall community performance. Although there have been other efforts using single-species probiotics to fight against pathogen colonization (*22*), this work represents a fairly simple microbial community as a way to suppress pathogens in the host intestine. Compared with the single strain system, in a community the individual strains appear to benefit from the complex mix of available nutrients and to acquire an enhanced ability to suppress organisms that are pathogenic to the host.

## Discussion

Gut microbe metabolism affects the health of animal hosts in numerous ways, from the liberation of indigestible nutrients (*4*) to the modification of host-secreted bile acids (*7*) to the synthesis of certain vitamins and neurotransmitters at significant levels (*1*). Moreover, metabolic interactions among members of the microbiota help maintain community structure, thereby preventing pathogen invasion or other dysbioses (*3*). *C. elegans* is a useful model host for microbiome studies, particularly in relation to development and host-pathogen interactions (*9, 10*). In this study, we developed a model of metabolic microbe-host interactions and directly demonstrated that colonization with heterologous bacteria enables *C. elegans* to digest and incorporate carbon from previously indigestible long-chain carbohydrates. In addition to isotopic measurement of carbon incorporation, we demonstrate direct benefits both to the host, with increased larval yield, and to other gut species, which in turn may provide protective effects against pathogens.

We developed assays to quantify the effects of an exogenously introduced microbiome on the nutritional status of *C. elegans.* Datasets collected towards this end include high-throughput *in vitro* carbon source growth screens to identify bacteria capable of degrading complex carbon sources and *in vitro* and *in vivo* screens to identify antibiotic cocktails for removal of colonized bacteria from the *C. elegans* gut. These results can support the use of these organisms as a convenient model for future microbiota-host interaction studies.

Even in these simple systems, good lessons have been learned about the intricacies of inter-species interactions in the potential pitfalls of designing a novel microbiome. First, as we have observed, interactions between microbes must be considered: the introduction of multiple functionally equivalent taxa (as with *P. cellulosa* and *B. subtilis*) does not necessarily increase the functional capacity of the community if there is exploitation competition between strains, which can reduce the functional redundancy that would otherwise be a desirable outcome of this colonization strategy. However, in our experiments, the use of a facultatively interacting taxon (*L. plantarum*) allowed us to add a new functional capacity (resilience against pathogen invasion) by capitalizing on the particular properties of this facultative species. This new function of the community was boosted when we exposed it to CMC, which can be utilized by cellulolytic *P. cellulosa* in the community, demonstrating that distinct bacteria can interact to combat pathogens via cellulose utilization. These results, consistent with other research (*23*) in this area, indicate that understanding interactions between bacterial strains within an *in vivo* host environment will be critically important in selecting novel colonists for a host microbial ecosystem. Community assembly starting from modular orthogonal interactions offers a promising approach for designing novel microbiome communities (*24*–*29*). Our results have suggested that nutritional interactions can enhance the performance of the system. Recent studies that utilize porphyran as an orthogonal metabolite to enable strain engraftment (*30, 31*) are consistent with this view.

More broadly, with respect to the specific results of these experiments, we believe that further efforts to expand the nutritional scope of animals could have a positive impact on food security. Globally, humans rely on a very slender thread of genetic diversity. In fact, according to the U.N.’s Food and Agriculture Organization (FAO), more than 50% of all human calories come from just three plants: rice, maize, and wheat (*32*). Engineering the gut microbiome could permit hosts to digest new food sources or enhance the efficiency of our current nutrition processing capabilities. With rapid progress in enzyme evolution (*33*) and microbiome engineering (*34*) techniques, facilitated by developments in model host organisms, engineering the gut microbiome could accelerate nutrient processing in humans or livestock or even expand diets to include entirely new food sources.

## Acknowledgments

The *C. elegans* native microbiome collection was a gift from the Dr. H. Schulenburg group, Christian-Albrechts University, Keil, Germany.

## Author contributions

Q.S., N.M.V., B.C., C.P.M., N.M., M.T., J.J.C., A.S.K., J.G., and T.K.L. designed and conceived the research. Q.S. designed and performed the bacterial CMC degradation capacity quantification assay. C.P.M designed and performed the bacterial CMC utilization assay. Q.S., N.M., and B.C. worked on the bacterial colonization and CMC degradation assay in the worm gut. B.C. conducted the ^14^C-cellulose utilization experiment. N.M.V. performed the worm *fat-7p::GFP* and larval yield experiments. M.T. and N.M.V. did the community experiment to suppress pathogens. C.P.M. helped with the final data analysis and figure processing.

