## Supplemental Material for "Colonization with heterologous bacteria reprograms a *Caenorhabditis elegans* nutritional phenotype"

Supplementary Materials:

Materials and Methods

*Bacterial species, worm strains, and culturing*

Bacterial species used in this study are listed in Table S1, along with species-specific growth media. In absence of specific culture requirements, bacterial cultures were routinely inoculated from glycerol stock and grown in LB media in a 30°C or 37°C shaking incubator or a benchtop shaker (23°C) for 24-48 hours before use.

Unless otherwise stated, N2 wild-type *C. elegans* (Caenorhabditis Genetic Center) were used in these experiments. Cultivation of all worms used here was performed at 25°C on NGM plates seeded with *E. coli* OP50. Synchronization of worms for these experiments was performed using standard protocols(*1*); briefly, worms were cultivated on NGM to obtain large numbers of gravid adults hermaphrodites, and eggs were isolated from these plates via bleach-NaOH lysis of adults. Eggs were rinsed 5-6X in M9 worm buffer to remove the hypochlorite solution and allowed to hatch out in 5 mL of M9 worm buffer overnight, after which the synchronized and starved L1s could be used for experiments. When reproductively sterile adult worms were desired, L1 larvae were plated onto NGM + 1 mM IPTG + 50 µg/mL plates seeded with *E. coli* expressing *pos-1* RNAi from a plasmid(*2*); worms grown to adulthood on these plates produce inviable eggs and are therefore reproductively sterile(*3*).

*Cellulolytic bacteria screening*

A Congo Red assay was used to determine cellulose hydrolyzing capability of bacteria. 1% CMC plates seeded with bacteria were cultured overnight at 30°C. The plates were then flooded with 0.1% Congo Red for 15-20min and then rinsed with 1M NaCl solution. Halo formed around the colony indicates cellulase indicates cellulase activity. For resting cell assays, cells in liquid culture were first washed once with buffer containing 50mM Tris-HCl (pH 8.0) and suspended in 1% CMC (carboxymethyl cellulose) from Sigma. Samples were collected after one hour and immediately mixed with 0.5mL of DNS reagents (10g/L dinitrosalicylic acid, 10g/L sodium hydroxide, 2g/L phenol, 0.5 g/L sodium sulfite). After incubation at 95°C for 10min, 1mL of 40% Rochelle salts was added to fix the color before measuring the absorbance at 575nm.

*Bacteria colonization and analysis*

Washed adult synchronized worms were resuspended in S medium and moved in 500µL aliquots to 15mL culture tubes with a final concentration of ~1,000 worms/mL. 500uL of bacterial suspension at 2X desired concentration was added to each culture tubes. Culture tubes were incubated with shaking at 300 RPM at 25°C. After 24 hours, colonized worms were washed 3 times with M9 worm buffer + 0.1% Triton X-100 and 2 times with M9 worm buffer to remove external bacteria. Worms were subsequently chilled to 4C for 15 minutes to stop peristalsis and treated for 10 minutes with a 1:2000 solution of commercial bleach to remove any remaining external bacteria, then washed 2X in M9 worm buffer + 0.1% Triton X-100 to remove bleach prior to disruption.

For manual disruption, worms were transferred to 3 mL M9 worm buffer + 1% Triton X-100 in a small (35 cm) petri dish (Fisher Scientific). Individual worms were pipetted out and transferred to 0.5 mL clear microtubes (Kimble Kontes) for manual disruption with a motorized pestle (Kimble Kontes Pellet Pestle with blue disposable pestle tips, Fisher Scientific, all digestions in 20 µL of buffer). Magnification was provided by a magnifying visor (Magni-Focuser Hands Free Binocular Magnifier 3.5X). After disruption, tubes were centrifuged 2 min at 12,000 RPM to collect all material, and the resulting pellet was resuspended in 180 μL M9 worm buffer (final volume 200 μL) before transfer to 96-well plates for serial dilution in 1X phospho-buffered saline (PBS). 10, 100 and 1000-fold dilution samples were plated on LB or ATCC Medium 2720 plates to check the colonization density.

*Colonized bacteria benefit from CMC*

Synchronized L1 worms from the N2 wild-type lineage were plated on solid agar on mixed lawns of *E. coli* OP50 and *P. cellulosa*, and allowed to grow to adulthood. Following this colonization, day 0 adult worms were separated into two aliquots. One went through disruption for colonization density as described above. The second aliquot was divided between wells of a 24-well plate with S medium, S medium with 0.1% glucose and S medium with 0.1% CMC (final volume 1 mL/well), covered with a BreatheEasy gas-permeable membrane, and incubated with shaking at 200 RPM at 25°C for 24 hours. Individual samples went through disruption and plating to check on nutrition’s effects on colonization density.

*Measurement of ^14^C-carbon incorporation*

Each gut colonization type was done in triplicate using 200 N2 synchronized worms. Samples were incubated at room temperature, with gentle mixing, in 1mL M9 Worm buffer containing 0.5% Carboxy-methyl cellulose and 1uCi/ml ^14^C-Uniformly labeled cellulose from *A. thaliana* for 72hrs. Antibiotic cocktails (100ug/ml ciprofloxacin and 500ug/ml carbenicillin) were provided at 48 hrs. Scintillation counts were performed after washing the worms with M9 buffer containing 0.1% TritonX. Washing of the worms was repeated until the scintillation counts in the wash supernatant reached background levels. Scintillation counts were performed using liquid scintillation analyzer (PerkinElmer Tri-Carb 2910 TR) in a high-efficiency LSC-cocktail (PerkinElmer Ultima Gold).

*Antibiotics cocktails for gut bacteria elimination*

High-throughput antibiotic susceptibility screen over our collection of soil bacteria was conducted with incubating bacteria antibiotics pooled and checked on bacteria density afterward. (Supplementary information) Once an effective antibiotic cocktail was identified, the best conditions to kill bacteria in the worm gut were determined. Worms were colonized with YFP-labeled *Pseudomonas citronellolis* (GmR) and treated with antibiotic cocktails (ciprofloxacin 100 μg/mL + carbenicillin 500 μg/mL). After 24 hours, fluorescent gut-associated colonies were checked; disruption and plating of intestinal contents will indicate if all live bacteria were cleared out from the worm intestine.

*Larval yield assay*

N2 wild-type worms were synchronized according to standard protocols(*16*), and L1 worms were transferred to 10 cm NGM plates containing OP50 only, OP50 + *P. cellulosa* (50 µL 10X bacterial suspension from 48 hour culture in 10 mL ATCC 2720, grown at room temperature with shaking), OP50 + *B. subtilis* (prepared identically to *P. cellulosa* using LB as growth medium), Worms were incubated on plates at 25°C for 46 hours, then washed off plates using M9 worm buffer + 0.1% Triton X-100, rinsed 3X to remove the bulk of extracellular bacteria, and picked as single worms into individual wells of a 384 well plate containing 25 µL S medium + 0.25X heat-killed OP50 (from 50X stock, cells concentrated 50X from stationary phase) + 25 µg/mL gentamycin (to prevent external growth of bacteria) ± carbon (glucose or carboxymethylcellulose). For each combination of bacteria and carbon source, 24 worms were assayed in each experiment. After 48 hours, larvae were counted in individual wells using 4X magnification and white light illumination on a Ti-E inverted microscope.

*Assay for nutritional status using an integrated fluorescent reporter*

We used transgenic worms expressing the *fat-7p*::GFP reporter fluorescent reporter as an indicator of host nutritional status(*4*). *fat-7p*::GFP worms were synchronized according to standard protocols and grown to day 0 adults on *pos-1* plates to prevent reproduction, then transferred to 6 cm NGM plates containing live *E. coli* OP50 *P. cellulosa, P. cellulosa* was grown at 25°C, in 5 mL ATCC 2720 then pelleted at 9000 RPM for 2 minutes and resuspended in S medium to concentrate the bacteria prior to seeding NGM plates with pre-existing OP50 lawns (50 µL of each suspension per plate). Worms were allowed to feed on these plates for 24 hours at 25°C to allow bacterial colonization.

Nutritional status was read on a BioSorter large object sorter using the 250 micron nozzle. Adult worms were transferred from plates to individual wells of a 24-well plate in S medium with heat-killed *E. coli* (to provide nitrogen and other nutrients) + gentamycin 10 µg/mL (to prevent growth of bacteria outside the worm) ± glucose or CMC at the indicated concentrations, covered with a BreatheEasy gas-permeable membrane, and incubated for 24 hours at 25°C with shaking at 200 RPM. After incubation, worms were washed to remove heat-killed bacteria, and total GFP fluorescence per worm was measured on the BioSorter; worms were gated based on extinction and time of flight (TOF) to isolate full-sized adult worms in these data, due to imperfect penetrance of the *pos-1* reproductively sterile phenotype in some experiments.

*High-throughput carbon source utilization screen*

18 species were inoculated from frozen glycerol stock into 2mL of their preferred rich medium (see Table S1), and grown in a shaking incubator at 30°C and 300rpm for 48 hours. Fast growing species exceeding OD ~1.0 in the first 24 h of this period were diluted 1:1000 into fresh rich media to grow for the remaining 24 h. All cultures were then centrifuged at 3500 rpm for 5 minutes, then resuspended in a M9 media formulation lacking any carbon source. OD600 was measured in a 96 well plate on a SpectraMax M5 platereader, and culture densities were normalized to OD 1.0. These cultures were diluted 1:20 into 96 well deep well plates containing 400uL of 12 different M9 media formulations, each with a different carbon source at 0.5% concentration (arabinose, cellobiose, carboxymethyl cellulose, ethanol, glucose, lactose, raffinose, soluble starch, sucrose, xylan hemicellulose, and xylose, as well as a no-carbon control). Cultures were grown at 30°C and 900rpm for 48 hours. OD600 was measured again, and the M9 no-carbon control cultures were used to subtract out apparent growth that may have occurred from residual rich media or absorbance changes independent of carbon utilization. Triplicate assays were performed on different days, with pairwise correlations of r^2^ ≥ 0.87 between days.

*Antibiotic susceptibility screen*

4 species were grown from frozen glycerol stocks for 48 hours in preferred rich media using the same protocol as in the carbon utilization screens. Cultures were centrifuged at 3500 rpm, and resuspended in fresh rich medium and normalized to an OD600 of 1.0 by diluting with additional rich medium. Normalized cultures were next diluted 1:100 into 96 well deep well plates containing 400uL rich media dosed with one of seven antibiotics (carbenicillin, chloramphenicol, ciprofloxacin, kanamycin, nalidixic acid, streptomycin, tetracycline, and an ethanol control) at 1x, 5x, and 10x working concentrations. Cultures were grown at 30°C and 900 rpm for 24 hours, then OD600 was measured on a SpectraMax M5 platereader to determine bacteriostatic potential. To determine whether each drug had proved bactericidal, five 5uL spots of each culture were grown on permissive nutrient agar. Colony counts were binned by order of magnitude (i.e. 0, 1-10, 11-100, lawn of >100 colonies).

*Engineered cellulolytic P. putida*

To engineer non-cellulolytic *P. putida* to cellulolytic bacteria. pET24a with ice-nucleation protein fusion with endoglucanase Cel5A assembled through Gibson assembly were transformed into *P. putida*. Induction of Cel5A surface display was achieved with 0.1 mM Isopropyl β-D-1-thiogalactopyranoside (IPTG) at OD 1 of bacteria culture. And overnight (16hr) induction at 16°C was conducted before bacteria was collected and reducing sugar assay from the bacteria was checked.

*Clearance of bacteria from the gut using antibiotics*

In order to drive clearance of bacteria from the gut, we screened antibiotics to develop a cocktail with efficacy against all species in our consortium. *In-vitro* tests identified a combination of 50 ug/mL carbenicillin and 10 ug/mL ciprofloxacin that should prove bactericidal to a broad range of species (Figure S5). However, during in-vivo experiments, we found that 10x higher concentrations of antibiotics were needed to effectively kill bacteria residing in the gut (Figure S5).

**Supplementary Figures**

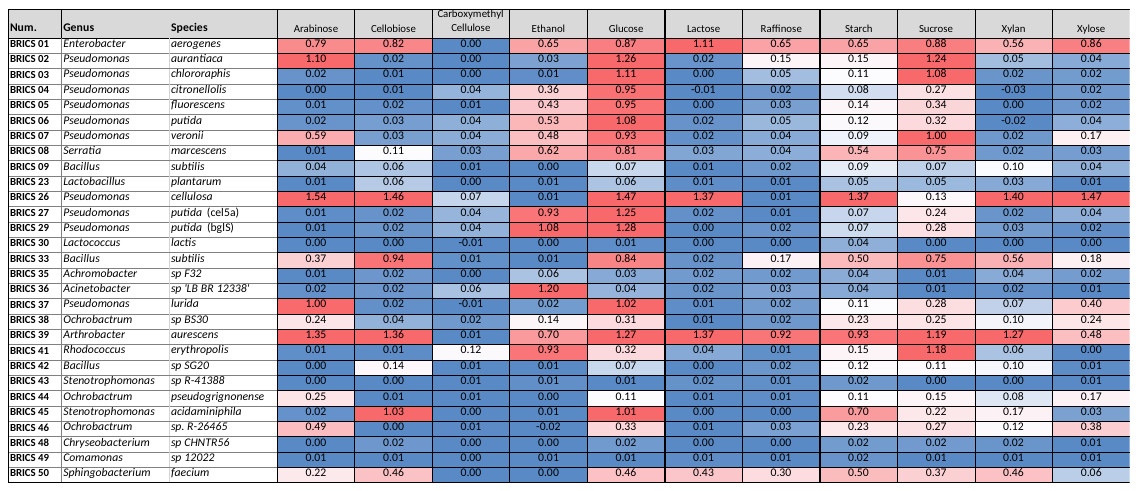

Figure S1. High-throughput *in vitro* carbon source growth screen. 29 species were tested (including *C. elegans* gut microbes) with 11 carbon sources in M9 minimal media. ΔOD600 vs. no carbon control over 48 hours agrees between 3 replicates: (R^2^ ≥ 0.87).

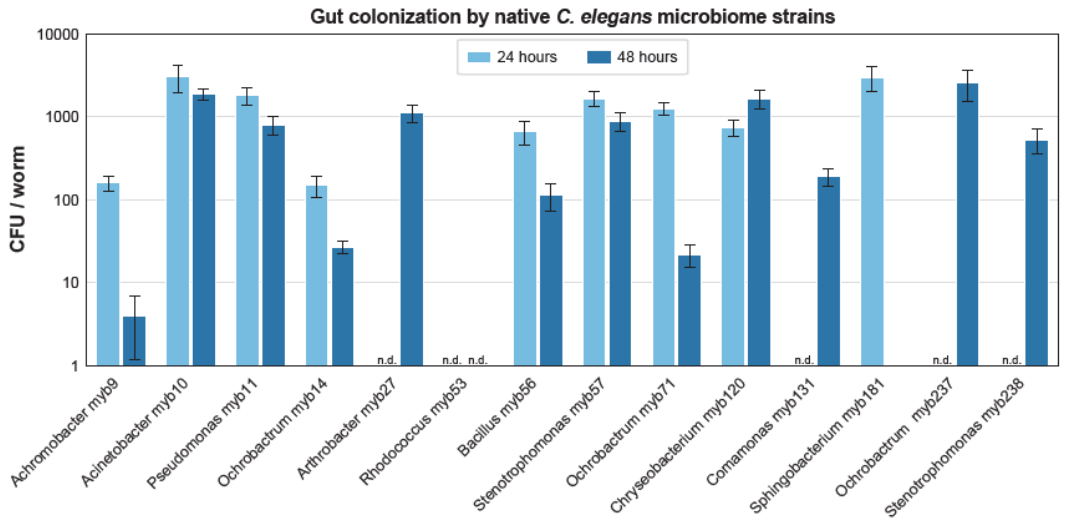

Figure S2. *C. elegans* native microbes colonize the worm gut. 16 native microbes (with strain names in Table S1) were each grown independently in LB media at 25C for 48 hours, diluted 1:10 from stationary phase, and used to colonize sterile N2 adult worms by feeding in liquid S medium at this concentration for 24 hours. After the initial feeding period, worms were removed from culture; a batch digest of 50 worms was performed (Time 0) to determine average CFU/worm for each microbial species, and the remaining worms were washed thoroughly and transferred to S medium + 1X heat-killed OP50 and incubated for a further 48 hours to determine persistence, after which a second batch digest (50 worms) was performed. Error bars represent mean ± SD of count error.

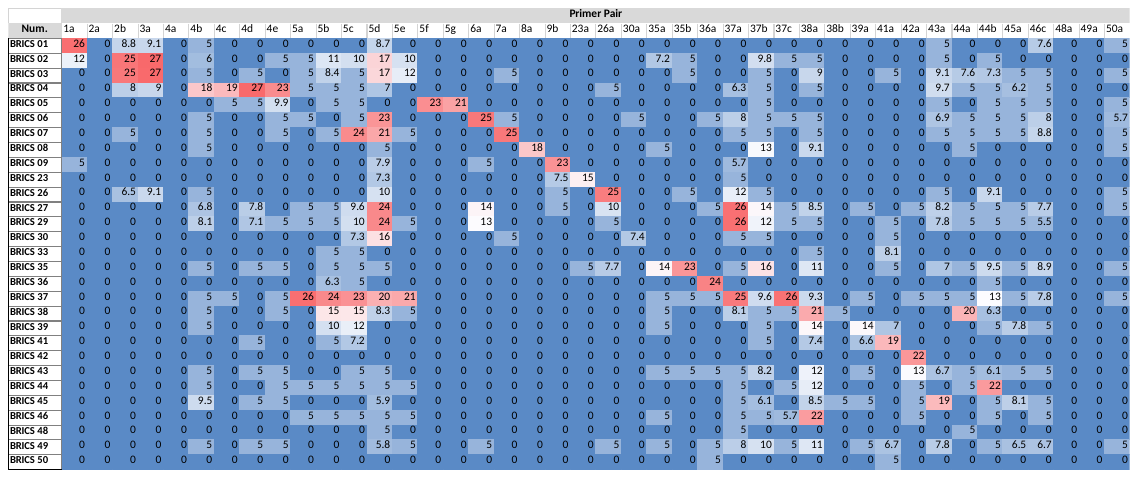

Figure S3. High-throughput screen or orthogonal qPCR identification primers for accurate identification of colonized bacteria. 29 species (including *C. elegans* gut microbes) were tested with 48 primers, with 1+ primer for each unique species, targeted to an annotated gyrB gene from closely related species. Displayed values indicate ΔCt compared to a no-template control signal that arose at approximately 35 cycles.

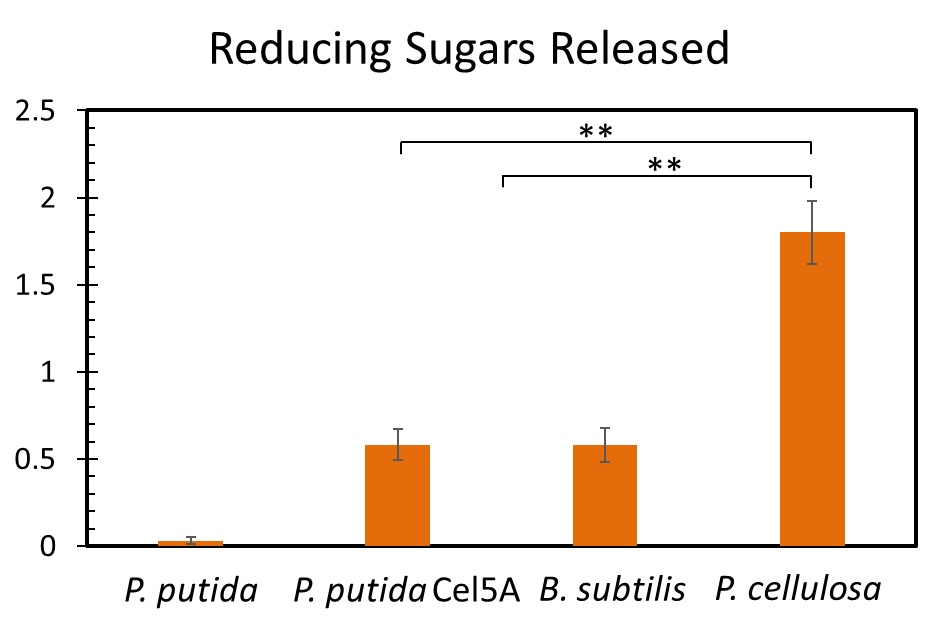

Figure S4. Quantification of cellulose-hydrolyzing capabilities of natural (*P. putida, B. subtilis, P. cellulosa*) and engineered (*P. putida* Cel5A) soil organisms. **P < 0.01, ***P < 0.001, Student’s t test. Error bars represent 95% confidence intervals of the mean.

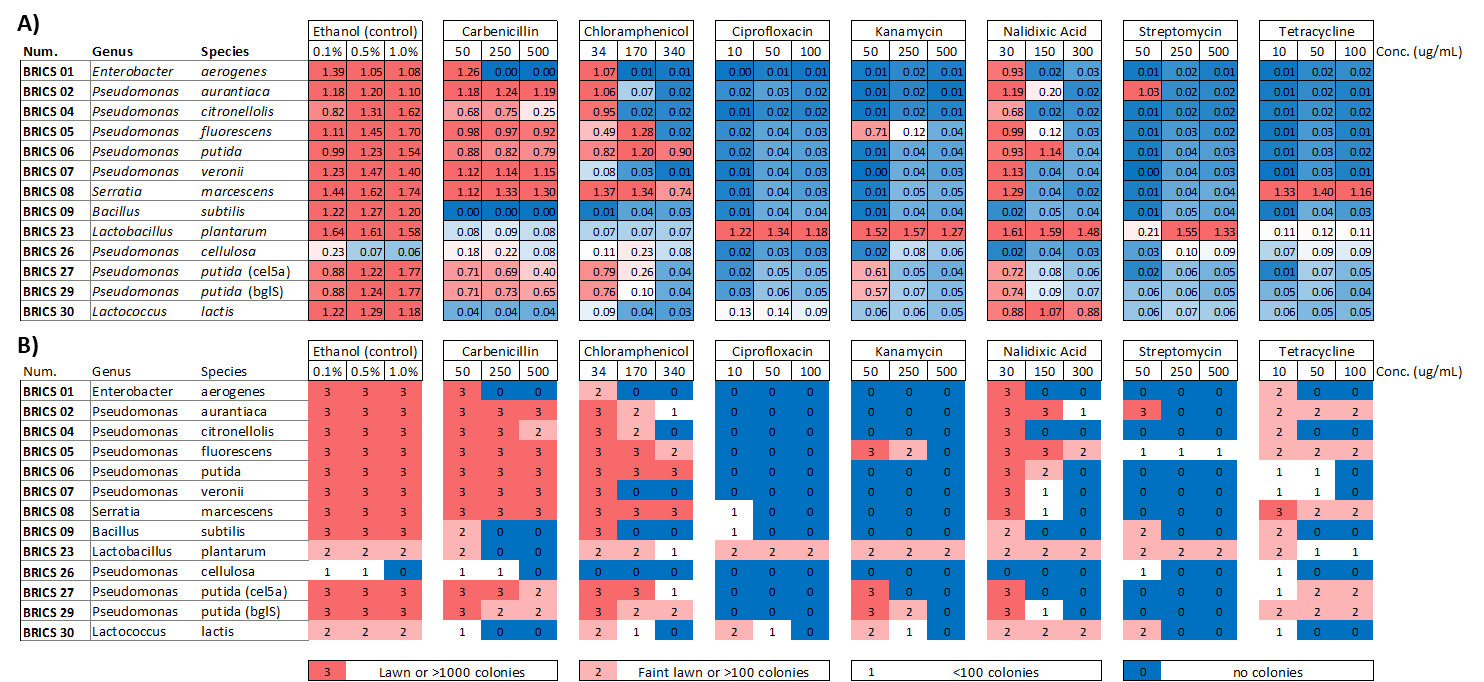

Figure S5. Large scale in vitro screen to identify antibiotic cocktail for bacteriostatic and bactericidal treatments. A) Optical density and B) colony counts were both checked to determine bacteriostatic and bactericidal potential, respectively.

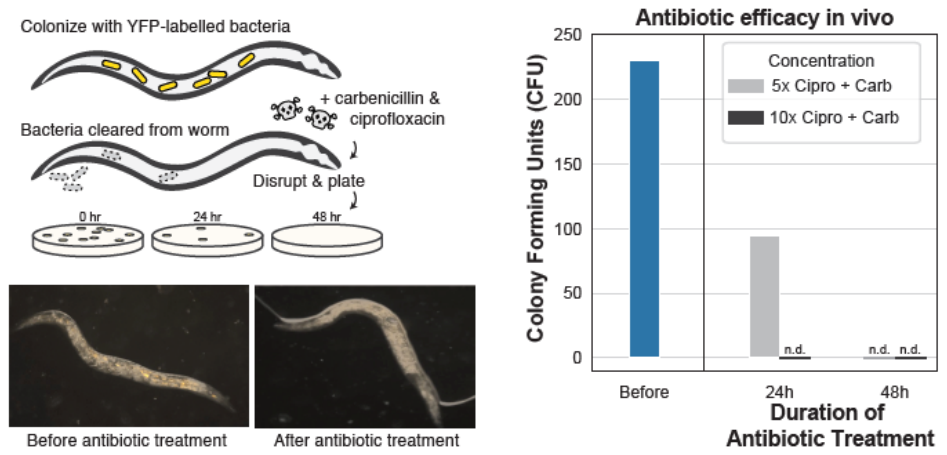

Figure S6. Plates assay (A) and microscope images (B) to validate removal of bacteria colonized in worm gut. For plates assay, *C. elegans* were disrupted and plated to check on colonized bacteria. For microscope images, *C. elegans* with YFP-expressing *P. citronellolis* before and after antibiotics treatment were checked under microscope for fluorescence bacteria in the gut.

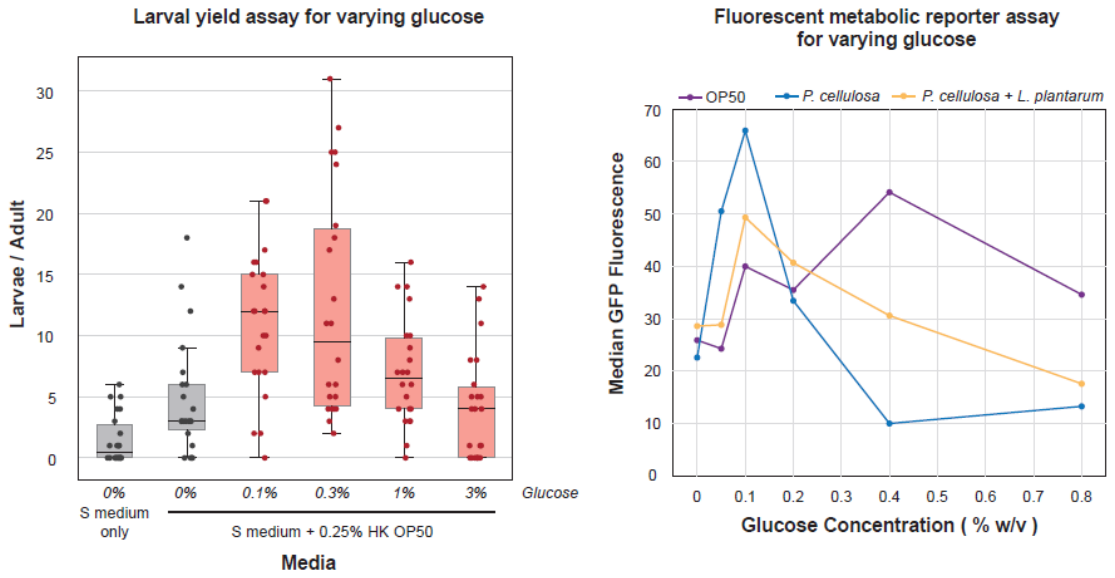

(B)

(A)

Figure S7. Optimal carbon source concentrations for nutritional benefit experiments. (A) In larval yield experiments, an optimum concentration of ~0.1%-0.3% glucose was determined. (B) Results from the *fat-7p*::GFP fluorescent reporter of host nutritional status (median GFP fluorescence per individual adult worm, ~1000 worms/condition) are consistent with the results of larval yield assays. It is plausible that the loss of benefit to the host at higher glucose concentrations is due to negative effects from osmolarity of the solution at these high concentrations.

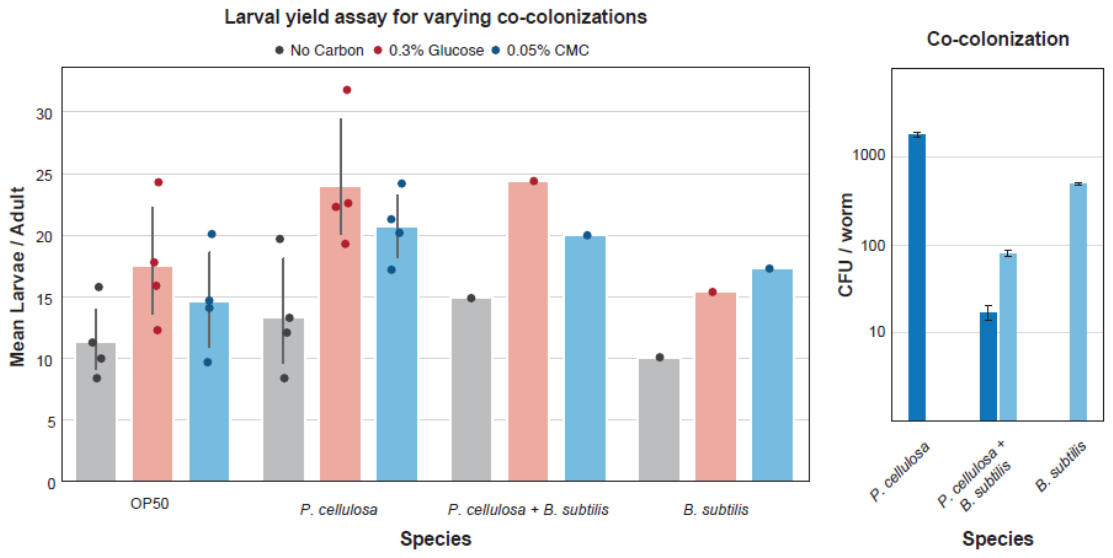

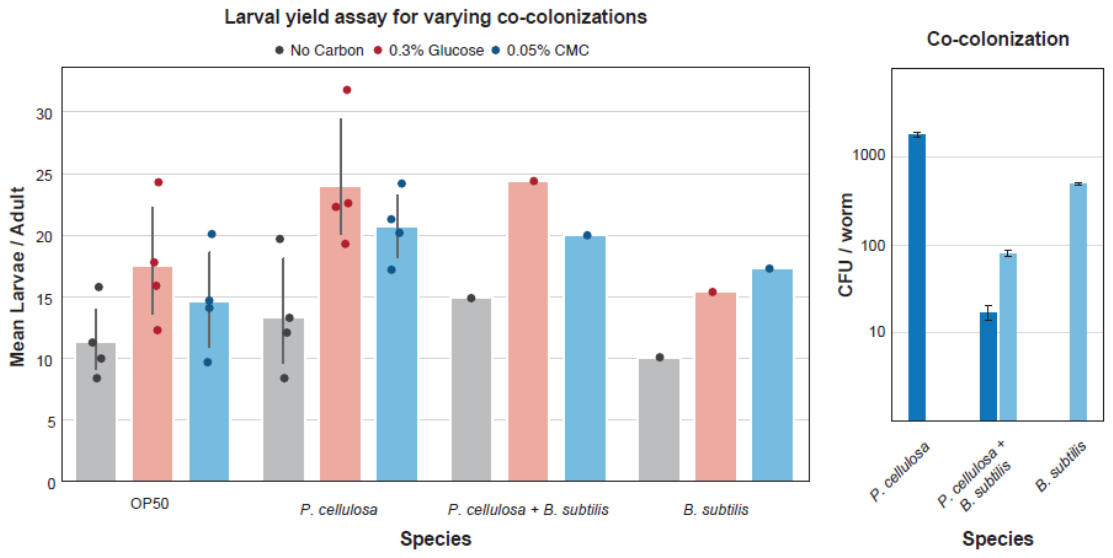

(B)

(A)

Figure S8. Co-colonization of the host with *Pseudomonas cellulosa* and *Bacillus subtilis* provides no additional nutritional benefit. (A) Co-colonization leads to reduced overall densities of bacteria in the gut, and particularly to suppression of *P. cellulosa*, indicating that this efficient cellulose degrader is out-competed by *B. subtilis*. (B) Consistent with these results, co-colonization does not increase the benefit of CMC supplementation to the host (wild-type N2 worms), as measured by larval output.

**Supplementary Tables**

| **Number** | **Genus** | **Species/Subspecies** | **Strain Designation** | **Source** | **Growth Media** |
| --- | --- | --- | --- | --- | --- |
| **BRICS 01** | *Enterobacter* | *aerogenes* | 13048 | ATCC | NB |
| **BRICS 02** | *Pseudomonas* | *aurantiaca* | 33663 | ATCC | NB |
| **BRICS 03** | *Pseudomonas* | *chlororaphis* | 9446 | ATCC | NB |
| **BRICS 04** | *Pseudomonas* | *citronellolis* | 13674 | ATCC | NB |
| **BRICS 05** | *Pseudomonas* | *fluorescens* | 13525 | ATCC | NB |
| **BRICS 06** | *Pseudomonas* | *putida* | 12633 | ATCC | NB |
| **BRICS 07** | *Pseudomonas* | *veronii* | 700474 | ATCC | NB |
| **BRICS 08** | *Serratia* | *marcescens* | 13880 | ATCC | NB |
| **BRICS 09** | *Bacillus* | *subtilis 168* | 23857 | ATCC | NB |
| **BRICS 23** | *Lactobacillus* | *plantarum* | 14917 | ATCC | MRS |
| **BRICS 26** | *Pseudomonas* | *cellulosa* | 55703 | ATCC | ATCC2720 |
| **BRICS 27** | *Pseudomonas* | *putida* | KT2440 (cel5a) | This Study | LB + Kanamycin |
| **BRICS 29** | *Pseudomonas* | *putida* | KT2440 (bglS) | This Study | LB + Kanamycin |
| **BRICS 30** | *Lactococcus* | *lactis subsp. Cremoris* | MG1363 | ATCC | GM17 |
| **BRICS 33** | *Bacillus* | *subtilis subsp. Spizizenii* | 6633 | ATCC | NB |
| **BRICS 35** | *Achromobacter* | *sp. F32* | myb9 | Dirksen et al 2016 | NB |
| **BRICS 36** | *Acinetobacter* | *sp. LB BR 12338* | myb10 | Dirksen et al 2016 | NB |
| **BRICS 37** | *Pseudomonas* | *lurida* | myb11 | Dirksen et al 2016 | NB |
| **BRICS 38** | *Ochrobactrum* | *sp. BS30* | myb14 | Dirksen et al 2016 | NB |
| **BRICS 39** | *Arthrobacter* | *aurescens* | myb27 | Dirksen et al 2016 | NB |
| **BRICS 41** | *Rhodococcus* | *erythropolis PR4* | myb53 | Dirksen et al 2016 | NB |
| **BRICS 42** | *Bacillus* | *sp. SG20* | myb56 | Dirksen et al 2016 | NB |
| **BRICS 43** | *Stenotrophomonas* | *sp. R-41388* | myb57 | Dirksen et al 2016 | NB |
| **BRICS 44** | *Ochrobactrum* | *pseudogrignonense* | myb237 | Dirksen et al 2016 | NB |
| **BRICS 45** | *Stenotrophomonas* | *acidaminiphila* | myb238 | Dirksen et al 2016 | NB |
| **BRICS 46** | *Ochrobactrum* | *sp. R-26465* | myb71 | Dirksen et al 2016 | NB |
| **BRICS 48** | *Chryseobacterium* | *sp. CHNTR56* | myb120 | Dirksen et al 2016 | NB |
| **BRICS 49** | *Comamonas* | *sp. 12022* | myb131 | Dirksen et al 2016 | NB |
| **BRICS 50** | *Sphingobacterium* | *faecium* | myb181 | Dirksen et al 2016 | NB |
| **NA** | *Escherichia* | *coli* | OP50 | Caenorhabditis Genetic Center | NB |

Table S1: Bacterial strains used in this study.

Table S2: qPCR Primers tested in this study.

| **Primer Num.** | **Sequence** | **Target Species** | **Pair** |
| --- | --- | --- | --- |
| pRCM224 | GTCACCCGCTGGGTTAATCA | 1 | a |
| pRCM225 | CTGCAGCTCGCTGTTTTCAC | 1 | a |
| pRCM226 | TCTGGGAACAGACCTACGTTCA | 2 | a |
| pRCM227 | GATGTTCTTGAAGGTCTCGCTGGA | 2 | a |
| pRCM228 | GGACAGTTCACGAATCCGCTT | 2 | b |
| pRCM229 | ACCCAGATTCACTTCAAGGCTTCTA | 2 | b |
| pRCM230 | ACGCAACCGTAAGACCCAG | 3 | a |
| pRCM231 | CCGACCTCTTGCGAGGAAAT | 3 | a |
| pRCM232 | TCTGGGAGCAGATCTATCGTC | 4 | a |
| pRCM233 | CCAGCTGAAGTGGATATTGGTAAA | 4 | a |
| pRCM249 | CTACAAGGTTTCCGGTGGCT | 4 | b |
| pRCM250 | GATCTGCTCCCAGACCTTGC | 4 | b |
| pRCM251 | ACAAGCTGGTCTCCTCCGA | 4 | c |
| pRCM252 | CTTGGCTTCGTTGGGGTTCT | 4 | c |
| pRCM255 | CGAAGGCAAGGTCTGGGAG | 4 | d |
| pRCM256 | GATGCTGGTGAAGGTCTCGT | 4 | d |
| pRCM257 | GAAGGCAAGGTCTGGGAGC | 4 | e |
| pRCM258 | TGCTGGTGAAGGTCTCGTTG | 4 | e |
| pRCM234 | ATGAAAATCGTTGGCGACAG | 5 | a |
| pRCM235 | CCGGAGTTGAGGAAGGACAG | 5 | a |
| pRCM243 | GAATCCACGGGTACGCAGAT | 5 | b |
| pRCM244 | CACCGGAGTTGAGGAAGGAC | 5 | b |
| pRCM245 | ATGGCGGTACTCACTTGGTG | 5 | c |
| pRCM246 | GGCGACTTTGTGCTTCTTGG | 5 | c |
| pRCM247 | ACTCACTTGGTGGGTTTCCG | 5 | d |
| pRCM248 | ATAATCGCGGTCAGGCCTTC | 5 | d |
| pRCM253 | GAATCCACGGGTACGCAGAT | 5 | e |
| pRCM254 | CACCGGAGTTGAGGAAGGAC | 5 | e |
| pRCM259 | AAAACCCTCAAGCGTCTTTCG | 5 | f |
| pRCM260 | AACGCCTTCAGCCAAATGTC | 5 | f |
| pRCM261 | GATAAGGCGCAGATGGACATT | 5 | g |
| pRCM262 | AACCATGGGAGGTCGTTTCA | 5 | g |
| pRCM236 | AAGTGGAAATCACCTCCCACG | 6 | a |
| pRCM237 | GTACGTAAGCACCTTCACCCAA | 6 | a |
| pRCM238 | TAAACGCCCTCTCCGAATTGC | 7 | a |
| pRCM239 | GCTTTCGCCAACAACCTTCATC | 7 | a |
| pRCM240 | GTGCACGAACAAACTTACAGCC | 8 | a |
| pRCM241 | GGTCACATTGGTAAAGGTCTGGT | 8 | a |
| pRCM307 | CGCCAAACCTATAAACGCGG | 9 | b |
| pRCM308 | TCAGGGTCCGGGACAAAATG | 9 | b |
| pRCM349 | GTGGGACGCATGAAGAAGGT | 23 | a |
| pRCM350 | TCTTCGCCAGATAGGTTCGC | 23 | a |
| pRCM311 | AGACAAGCTGGTGTCTTCCG | 26 | a |
| pRCM312 | CCACTGCCTTGGCATCATTG | 26 | a |
| pRCM351 | GCAGACGAATATGACGCCAG | 30 | a |
| pRCM352 | GGTGCAACCCTTCTTTTGAGG | 30 | a |
| prCM407 | AGTTCAGCAGCCAGACCAAG | 35 | a |
| prCM408 | TTCTCGAGCAGCCAGGATTC | 35 | a |
| prCM415 | GAACAACGGCGTCAAGATCC | 35 | b |
| prCM416 | TTGGCGCGGTTGATGTATTC | 35 | b |
| prCM440 | GCGATTGAAACAACGCAACC | 36 | a |
| prCM441 | CGCGGGAAATAAATCGTCGT | 36 | a |
| prCM399 | CCGAAGGTCTGGCCAAGAAG | 37 | a |
| prCM400 | TCCGGTACCTTCACCGAGAT | 37 | a |
| prCM403 | ATCGTTACGACCGCAACCTG | 37 | b |
| prCM404 | AGTGCATCCAGCCCTTCG | 37 | b |
| prCM401 | AAGGTCTGGCCAAGAAGCAT | 37 | c |
| prCM402 | GATCCGGTACCTTCACCGAG | 37 | c |
| prCM397 | AGACCATATGATGCAGGCCG | 38 | a |
| prCM398 | CGCGGATTCCATTCGTGTTC | 38 | a |
| prCM442 | CTTTGCGACGAAGTACAGGC | 38 | b |
| prCM443 | TATGGCACTGCTCGGTTCTG | 38 | b |
| prCM448 | GCTTTCGAAACCGAGGACAC | 39 | a |
| prCM449 | GTGTTTGCGTACGTGTGGAC | 39 | a |
| prCM450 | GCTCCAACGATCCGGAGAAG | 41 | a |
| prCM451 | CTGGTACATCGAGTCTCGGC | 41 | a |
| prCM446 | ACACCATGCAAACCACCAGA | 42 | a |
| prCM447 | ATGGGACGTCCTGCTGTAGA | 42 | a |
| prCM454 | TGCTGGTGGATGTGTTCCAG | 43 | a |
| prCM455 | GTTTGGTGGTGGTTTCCAGC | 43 | a |
| prCM464 | AGGACAAGCTCGTTTCCTCG | 44 | a |
| prCM465 | TTCGACGATGGTCTTGCCAC | 44 | a |
| prCM466 | TCAAGCTTACAGACCGTCGC | 44 | b |
| prCM467 | AGCGACTTCTTGCTCTGGTC | 44 | b |
| prCM468 | GCTGATGGAGGAAGCCAAGA | 45 | a |
| prCM469 | GTATCGGGGTTGACCGTGG | 45 | a |
| prCM452 | AAGAAGCCACGCTGGAAATG | 46 | c |
| prCM453 | CGTGTGAAGACCCGCCAG | 46 | c |
| prCM458 | CCGACGAAAGAGAAAGGTTGG | 48 | a |
| prCM459 | TGGATTCCCTGCTTCCATCG | 48 | a |
| prCM460 | GTCAGCTCCGAAGTCCGTG | 49 | a |
| prCM461 | TCGACAATCTTGTTGCAGAGAATC | 49 | a |
| prCM462 | TGGTGTAACCCTTCGCCAC | 50 | a |
| prCM463 | CGTGCGCAAGAATATGACGC | 50 | a |
